# Diversity of entomopathogens Fungi: Which groups conquered the insect body?

**DOI:** 10.1101/003756

**Authors:** João P. M. Araújo, David P. Hughes

## Abstract

The entomopathogenic Fungi comprise a wide range of ecologically diverse species. This group of parasites can be found distributed among all fungal phyla and as well as among the ecologically similar but phylogenetically distinct Oomycetes or water molds, that belong to a different kingdom (Stramenopila). As a group, the entomopathogenic fungi and water molds parasitize a wide range of insect hosts from aquatic larvae in streams to adult insects of high canopy tropical forests. Their hosts are spread among 18 orders of insects, in all developmental stages such as: eggs, larvae, pupae, nymphs and adults exhibiting completely different ecologies. Such assortment of niches has resulted in these parasites evolving a considerable morphological diversity, resulting in enormous biodiversity, much of which remains unknown. Here we gather together a huge amount of records of these entomopathogens to comparing and describe both their morphologies and ecological traits. These findings highlight a wide range of adaptations that evolved following the evolutionary transition to infecting the most diverse and widespread animals on Earth, the insects.

## Introduction

The Kingdom Fungi is one of the major groups of eukaryotic microbes in terrestrial ecosystems (Mueller and Schmit 2007). There are approximately 100,000 described species of Fungi (Kirk *et al.* 2008), which only represents a fraction of its diversity, estimated to be between 1.5 and 5 million species (Hawksworth and Rossman 1997, Blackwell 2011). Importantly, one of the hallmarks of Fungi is their propensity to form intimate interactions/associations with other groups of life on Earth (Vega and Blackwell 2005). According to Hawksworth (1998), 21% of all species of described Fungi are associated with algae as lichens and 8% form intimate relationship with plants as mycorrhiza, being this a prominent example of such close association, which occur in rhizosphere (around roots), where arbuscular mycorrhizal and ectomycorrhizal fungi form associations with plants. Few if any organisms in terrestrial ecosystems exist in nature in the complete absence of Fungi and for this reason they are essential players in the maintenance of ecosystem health. Another group of importance are the Oomycetes. These are so-called water molds and belong to a very distant Kingdom (Stramenopila), more closely related to brown algae (Kamoun, 2003). However, it is appropriate to discuss them with fungi as they were long considered to be fungi and are ecologically very similar.

The Insects with over 900,000 described species represent the most species-richness groups of eukaryotes (Grimaldi & Engel, 2005, pg. 12). They are known to form intimate relationships with many fungal groups: mutualistic endosymbionts that assist in nutrition (Suh et al. 2005), fungi as food sources that insects farm (Currie et al. 2003; Mueller *et al.* 2005), sexually-transmitted parasites and commensals (DeKesel 1996), and pathogens suggested to have pronounced effects on host populations (Evans & Samson 1982; 1984). However, even though we know many different fungal-insect associations exist this area remains one of the most understudied areas of fungal biodiversity and likely harbors one of the largest reservoirs of undocumented fungal species (Vega and Blackwell 2005).

A prominent characteristic of insects is a chitinous exoskeleton, which the great majority of entomopathogenic fungi and oomycetes need to penetrate and overcome the cellular and humoral defenses in the hemocoel (Evans, 1988). Following entry some groups (i.e. *Metarhizium* and *Beauveria* in the Order Hypocrelean, Phylum Ascomycota) grow inside the host as yeast-like hyphal bodies, multiplying by budding (Prasertphon & Tanada 1968). However, some groups grow differently, for instance in some Entomophthoraceous fungi which produces protoplasts (cells without cell walls) or in the case of entomopathogenic Oomycetes, Chytridiomycetes and *Entomophthora aphids*, which grows as hyphae inside host’s body (Roberts & Humber, 1981; Zattau & McInnis, 1987; Lucarotti & Soulkamy, 2000). After the colonization of the insect’s body, the fungi and water molds need to grow structures to produce and spread their spores. The majority of entomopathogenic fungi kill their hosts before spore production starts, but few of them, especially some Zygomycota species, sporulate from the living body of their hosts (Roy et al. 2006, see this study). All of entomopathogenic water molds kill their insect hosts as a developmental necessity

All entomopathogenic fungi and water molds transmit via spores. There are two spore types to consider. The sexual meiotic spores (zoospores, zygospore, basidiospore, ascospore) are actively released into the environment and asexual mitotic spores (conidia) passively released (Roberts & Humber, 1981). Each group of these entomopathogenic organisms presents their particular strategy and morphology that makes them very efficient pathogens, as will be discussed further for each group. Having provided a short introduction, the aim of this paper is to ask which groups of Fungi and Oomycetes evolved the ability to exploit the insect body. Before doing that we first provide with overall characteristics of the fungal/water mold groups that contain taxa that infect insects since many groups are generally unfamiliar.

## Oomycota

The species belonging to Oomycota were in the past, traditionally called also as fungi. However, phylogenetic studies confirmed that these organisms do not share a common ancestral with fungi, being these allocated in the Straminipila (Alexopoulos, 1996), a kingdom that present morphologically diverse organisms such as Hyphochytriomycota and Labyrinthulomycota (Alexopoulos, 1996). In addition, the phylum Oomycota presents a number of biological and morphological characters that can distinguish them from Fungi. The first one is the asexual reproduction by biflagellate zoospore with a longer tinsel flagellum directed forward and a shorter whiplash flagellum directed backward (Barr, 1992; Dick, 2001). Regarding the sexual reproduction, they have a unique oogomous reproduction by gametangial contact, resulting in a thick-walled sexual oospore. At a cellular level, they possess mitochondria with tubular cristae, whereas the fungi have mitochondria with flattened, plate-like cristae. Also, and distinctly, their cell walls contain cellulose whereas the cell walls of fungi contain chitin (Alexopoulos, 1996).

This group of microorganisms has evolved both pathogenic and saprophytic lifestyles (Phillips *et al.* 2007). As a pathogen, oomycetes are able to infect a broad range of host such as algae, plants, protists, fungi, arthropods and also vertebrate animals and humans, being able to lead them to death (Kamoun, 2003). Certain genera are well-known plant pathogens, such as *Phytophthora* that is responsible for the Irish Potato Famine (Haas *et al.* 2009) and sudden Oak death (Brasier & Weber, 2010). Those taxa infecting arthropods (the group containing insects) have been recorded from lobsters (Fisher *et al.* 1975; Khulbe, 1994) and shrimps (Hatai *et al.* 1992) as well as insects (Seymour, 1984; Samson *et al.* 1988; Pelizza *et al.* 2009; Stephen & Kurtböke 2011). As an insect parasite, they are known to infect the mosquito’s genera *Aedes, Anopheles* and *Culex*, all vectors of human diseases.

## Microsporidia

For long time the Microsporidia’s species were classified within the Phylum Apicomplexa as “sporozoans parasites” (Margulis & Schwartz, 1998). However, an increasing number of studies have strongly suggesting that Microsporidia are within the Kingdom Fungi (Hirt *et al.* 1999; Keeling *et al.* 2000; Keeling 2003; James *et al.* 2006; Hibbett *et al*. 2007). Although this is not resolved yet, since some of these studies placed the Microsporidia as sister group of *Rozella allomycis*, a chytrid (James *et al.* 2006), and others placed them as close related to Zygomycota (Keeling *et al.* 2000). However, a conclusive resolution about Microsporidians among the Fungi will require further genetic studies from basal fungal taxa (James *et al.* 2006). Nevertheless, based on the studies mentioned above and the Microsporidia’s ecological function as insect pathogens we will include them among the Fungi on this study.

The most remarkable feature of this group is its unique spore that acts as telescopic syringe injecting its protoplast material into the host. This structure is responsible for performing the transmission between their hosts and it is the only viable stage outside the host’s cell (Keeling & Fast, 2002). The spore is very distinct, being relatively small, ranging from 1 to 40 μm (Vávra & Larsson, 1999 pg.8), thick walled and containing a characteristic polar thread-like tube apparatus (Wittner, 1999). Oshima (1937), suggested that the protoplasm is transmitted from the Microsporidian spore to inside the host cell through the hollow tube formed during adherence and the subsequent discharge of the parasite’s intracellular content within the host’s cell. The discharging of polar tube occurs by breaking through the thinnest region of the spore wall, the apex. This event is compared by Keeling and Fast (2002) as “*likened to turning the finger of a glove inside-out*”.

The host range for most Microsporidian species is usually restricted, although some species with a wide host range have been described (Wittner, 1999 pg.2). They have been reported infecting a great number of domestic and wild mammalian species (Snowden & Shadduck, 1999, pg.393) fish (Shaw & Kent, 1999, pg.418), avian (Kemp & Kluge, 1975), amphibian, reptiles and even human immunocompromised AIDS patients (Didier & Bessinger, 1999, pg.225). For further informations see: Becnel & Andreadis, 1999; Sokolova *et al.* 2006; Briano *et al.* 2006; Lange, 2010; Kyei-Poku *et al.* 2011; Hossain *et al.* 2012; Vega & Kaya, 2012.

## Chytridiomycota

The phylum is suggested to be the earliest diverging lineage of the Fungi (James *et al.* 2006). There are reports of them dating from Lower Devonian (Taylor *et al.* 1992) and a parasitic chytrid-like Fungi dating from Permian on the Antarctica continent (Massini, 2007). The Chytrids (as they are traditionally called) are the only group of the Kingdom Fungi that possess motile cells at least once in its lifecycle. These cells are called zoospores, equipped with a single, posteriorly directed whiplash flagellum, which reflects their aquatic lifecycle (for details see Barr & Désaulniers, 1987). These zoospores respond to chemical gradients allowing them to actively locate their hosts, especially important for species pathogenic on aquatic organisms. Another important characteristic of the zoospores is the ability to encyst when the environment turns to cold, hot or dry, avoiding water loss or cell’s collapse (Gleason & Lilje, 2009). As mentioned before, the Oomycota also have motile spores and so this is an example of convergent evolution as both groups are aquatic.

The majority of chytrids are found as saprophytic organisms especially in fresh waters and wet soils, while there are some marine species (Thomas *et al.* 2012). Although, a significant number of species are known to be parasites of plants, animals, rotifers, tardigrades, protists and also other fungi (Sparrow, 1970; Martin, 1978; Karling, 1981; Dewel, 1985). Despite infections caused by Fungi (especially Asco- and Zygomycota) in insects are common (Evans & Samson, 1982), diseases caused particularly by Chytrids appear to be comparatively rare (Karling, 1948). For further reading see: (Voos, 1969; Whisler *et al.* 1975; Millay & Taylor, 1978; Kerwin, 1983; Couch & Bland, 1985; Padua *et al.* 1986; Powell, 1993).

## Zygomycota

The Zygomycota was traditionally organized as a single phylum in two classes, Zygomycetes and Trichomycetes. Both, sharing characteristics such as coenocytic mycelium (i.e. lacking regular septation), asexual reproduction usually by sporangiospores and absence of flagellate cells and centrioles (Alexopoulos, 1996). Their main general characteristic is the production of thick walled resting spore (i.e. zygospore) within a commonly ornamented zygosporangium, formed after fusion of two specialized hyphae called gametangia (Alexopoulos 1996). But, since they are not a monophyletic group, some morphological features may vary.

Nevertheless, after phylogenetic studies, this group was reorganized in five new taxa: Glomeromycota (for arbuscular mycorrhyzal Fungi), and all other zygomycetes distributed in five subphyla (Entomophthoromycotina, Kickxellomycotina, Mucoromycotina and Zoopagamycotina) without assignment to any phylum (James *et al.* 2006; Hibbett *et al.* 2007). Thereafter, Humber (2012) proposed a new monophyletic phylum, the Entomophthoromycota, with around 180 species, primarily obligate arthropod parasites and so-called entomophthoraceous fungi. (see the phylogeny of the group in Gryganskyi *et al.* 2012).

They are ecologically very diverse, widely distributed and very common. Most species are saprotrophic in soil and dung, where some of them are fast growing and can often be found on mouldy bread, fruits and vegetable (Moore et al. 2011). On the other hand, another groups are known to be parasites. Some are obligate mycoparasites (Dimargaritales), though another parasites of mushrooms can also be found in other groups (Zoopagales and Mucorales). In addition, some groups are known to be parasites of amoeba, rotifers and small animals, where the insect pathogens can be highlighted due their diversity (Moore *et al.* 2011).

The trichomycetes (now placed among the Mucoromycotina within the order Harpellales – Hibbett *et al.* 2007) are fungi, which exclusively inhabit the guts of various arthropods (Horn & Lichtwardt, 1981). However, since the trichomycetes apparently do little, if any harm to their hosts under natural conditions (Horn & Lichtwardt, 1981) and the nature of their relationships is not fully understood, they were not included in this study.

## Basidiomycota

This group, together with Ascomycetes, forms what are so-called superior fungi. They host some of the most well known fungi such as mushrooms, boletes, puffballs, earthstars, smuts, rust fungi, etc. They can be characterized by the formation of sexual spores (i.e. basidiospores) outside specialized reproductive cells called basidium. These spores in most cases are actively released, in other words, discharged forcibly by elaborate discharge mechanisms (Webster *et al.* 1984; Pringle *et al.* 2005). Another important and exclusive trait for the group is the clamp connections. These are structures formed during the division of the nuclei in the tip of growing hyphae, which help to ensure the dikaryotic condition (i.e. two nuclei in the same cell) (Alexopoulos 1996) and can also be used to identify the Phylum members, even in fossil records (Krings *et al.* 2011).

The Basidiomycetes, as they are commonly called, present important ecological traits. They occur colonizing dead woods, decaying cellulose and lignin as well as leaf litter decomposers on the forest floor (Braga-Neto *et al.* 2008), being essential components of forest ecosystems (Alexopoulos 1996). As pathogens, they present species infecting mainly plants (so-called smut and rusts) being responsible for huge losses in agriculture. Also forest environments are attacked by Basidiomycetes species like *Armillariella mellea*, which attack trees and *Heterobasidion annosum*, which attack conifers (Kendrick, 2000). As animal pathogens some species are known to attack nematodes, such as the anamorph phase (*Nematoctonus*) of the mushroom *Hohenbuehelia* and *Resupinatus* (Basidiomycota – Agaricales). Another interesting case occurs in the widely cultivated genus *Pleurotus*. Since they are primary colonizers of dead wood, which is deficient in nitrogen, the nematodes seems to be an important component of their diet. They exploit the nematodes by secreting substances that are able to inactivate them, allowing the fungus to colonize their sluggish bodies (Kendrick, 2000). As insect pathogens, few genera are known, which infect mainly scale insects (i.e. *Septobasidium* and *Uredinella* - Septobasidiales) and termite eggs (i.e. *Fibulorhizoctonia* - Atheliales).

The Septobasidiales attack exclusively scale insects (Hemiptera – Diaspididae (Evans, 1989)). The order includes two genera of entomopathogens, *Uredinella*, that attack just single insects and *Septobasidium*, attacking whole colonies with up to 250 individuals. This character is one of the most remarkable differences between both genera, but also the fungus size and micro-morphological features can be used to separate them. For example, the presence of a different type of spore in *Uredinella*, called by Couch (1937) as binucleate uredospore, not found in *Septobasidium*. In this same study, where the author describes the new genus *Uredinella*, he called it as “A new fungus intermediate between the rusts and *Septobasidium*” containing traits of both.

Another group within Basidiomycota was described as an opportunistic pathogen of termite eggs (Matsuura *et al.* 2000). This fungus was found living inside termite’s nest, among their eggs and were proven to act as pathogen of them in few occasions. The authors classified this fungus within Atheliales, as being a species of *Fibulorhizoctonia* (anamorph of *Athelia*), based in molecular studies (Matsuura *et al.* 2000).

## Ascomycota

The Phylum is the largest group in Kingdom Fungi. The majority of species are filamentous, producing regularly septate hyphae. They are characterized by the formation of sexual spores (i.e. ascospores) in sac-like structures called ascus. They are very diverse regarding the ecology, presenting members that are decomposers, plant pathogens, human and animal pathogens etc. In addition, the great majority of lichenized fungi belong to Ascomycota (Moore *et al.* 2011). As entomopathogens they are also the larger group among Fungi, attacking a wide range of insects (Figure 1).

**Figure 1:**
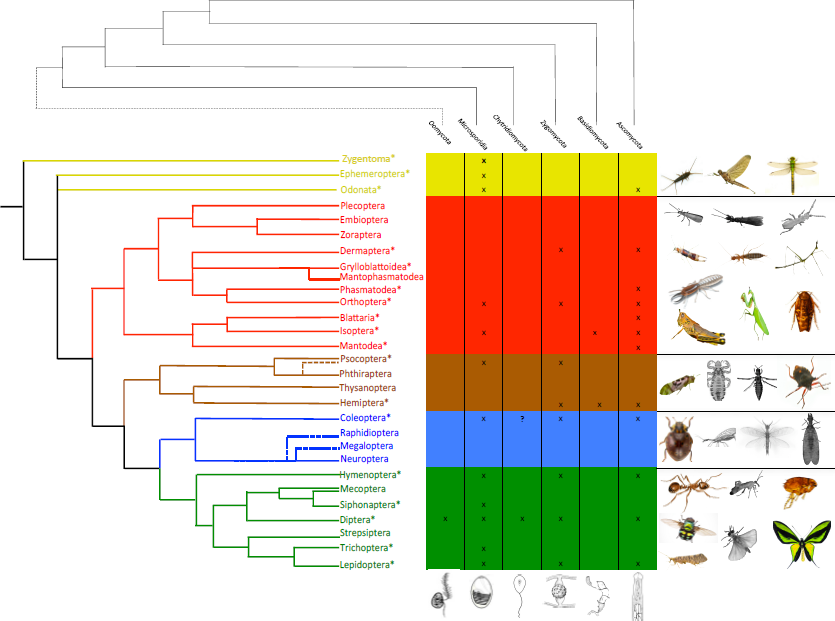
Insect orders x Fungal Phylla: The parasitic relationship between entomopathogens Fungi and their hosts. On the left, the phylogeny of insect orders (Grimaldi & Engel, 2005), on the top the phylogeny of fungal phyla and Oomycetes (adapted from James et al 2006), the table showing which fungal group infects each insect order. The colored insect figures means that the groups have fungal infection, the ones in gray means no recorded to have fungus as pathogen.

In the great majority of species they present a well-developed parasitic phase that infects the host’s body, before killing them. Furthermore, after killing the insect, this group is also able to colonize the cadaver saprophytically, keeping the hyphae growing even after the host’s death and them growing fruit bodies (Evans, 1988). According to the same author, these entomopathogens would have evolved at an early stage in moist tropical forests, particularly rainforests, attacking a wide range of different hosts since then (figure 1). The great majority of entomopathogens Ascomycetes form their spores inside structures called perithecia, a subglobose or flask-like ostiolate ascoma (Kirk et al. 2001), presenting a great diversity of spore types and shapes (Alexopoulos, 1996). The Phylum comprises from small groups of insect pathogens such as Pleosporales, Myriangiales and Ascosphaerales to the biggest group of entomopathogens, the Hypocreales (Samson et al. 1988).

Within the Pleosporales, the entomopathogenic species belong to the genus *Podonectria* (Petch, 1921), presenting unusual aspects among its class, like color and fleshiness. All known species were found infecting scale insects, covering the whole surface of insect body with cotton-like crust on which the perithecia will be produced and later, the multi-septated spores, not disarticulating into part-spores (Kobayasi & Shimizu, 1977). The genus is wordwide distributed, especially in warm climates (Petch, 1921) and the anamorphs related are the sporodochial genus *Tetracrium* (Petch, 1921; Kobayasi & Shimizu, 1977) and the pycnidial genus *Tetranacrium* (Rossman, 1978; Roberts & Humber 1981).

The Myriangiales includes a number of species associated with plants, resins, or scale insects on plants (Alexopoulos, 1996). The entomopathogen species present perennial growth for several years, or at least until the branch, where the scale insect is attached, dies. The hosts can be found directly under each stroma, where there are several scales, penetrated and covered by mycelium (Miller, 1940) or sometimes can be formed at the side of the insect (Petch, 1924). The growth occurs when the rains begin, where the live parts of the fungus body increase in diameter, producing ascocarps and later on ascospores. Reproduction is entirely by ascospores and no evidence of conidial (asexual spore producing cells) formation was found on stromata of any age neither in culture (Miller, 1938). For taxonomic and additional discussions see Petch (1924) and Miller (1938; 1940).

A unique group of bee pathogens are also included in this Phylum. The genus *Ascosphaera* is a group of approx. 26 species, which are specialists in exploit bee’s provisions. They are mostly exclusively saprophytes on honey, cocoons, larval feces or nest materials such as leaf, mud or wax of bees (Wynns et al. 2012). However, some species are known as widespread fungal disease agents, attacking broods of numerous species of solitary and social bees, in a disease called chalkbrood (Klinger et al. 2013). The infection occurs when the larva ingest fungal spores, being killed by the hyphal growth inside its body. Then, leading to new spore formation on the cuticle of the dead larvae (Vojvodic et al. 2012). The morphology of *Ascosphaera* is very peculiar when compared to other fungal groups. The ascoma is a small brown to blackish brown spore cyst, a single enlarged cell containing ascospores (Wynns et al. 2012). In addition, they also lack a well-developed ascocarp, which led them to be considered as yeasts (Alexopoulos, 1996). For a detailed life cycle see McManus & Youssef (1984).

The Hypocrealean fungus encompasses important genera of entomopathogens Fungi such as *Hypocrella*, *Cordyceps*, *Ophiocordyceps*, *Moelleriela* among others. In addition to these species, there are a myriad of anamorphic species related to them, i.e. *Hirsutella*, *Metarhizium*, *Hymenostilbe*, *Akanthomyces*, etc (Roberts & Humber, 1981). In the past, these anamorphic species (which just asexual stage is known) were treated traditionally as a separate group, the Deuteromycetes. But, nowadays with the advancing molecular studies, some of these species are now strongly supported to be asexual stages of mainly Ascomycota (see Liu et al. 2000). Even 25 years ago, Evans (1988 book) included both in the same section in his book’s chapter and said: “*The entomopathogenic Deuteromycotina are included within this subdivision* (Ascomycotina section of his book) *since many of them are either proven or purpoted anamorphs of Ascomycotina*”. Here, since the great majority of anamorphic species were proven to be part of life cycle of ascomycete teleomorph phase, we will just address the teleomorph names to avoid some synonymy and to follow the new “One fungus one name” code.

## Methods

To understand the current diversity of entomopathogenic fungi and Oomycetes, we gathered and analized a huge amount of papers and books published from 1785 to 2013. These materials were combined then with a search in specific websites about fungal diversity. A valuable initial source was the http://cordyceps.us. an electronic monograph of Cordyceps and related Fungi, where the visitor can find a great number of literature and species descriptions. We also consulted http://apps.webofknowledge.com. a useful website for journals search. To check the species names and its status we used the standard website for fungal searches: http://www.speciesfungorum.org and http://www.mycobank.org. We also counted with the valuable help of many specialists such as Dr. Richard Humber (USDA), Dr. Harry Evans (CABI), Dr. Bushan Shrestha (National Academy of Agricultural Science – South Korea). In our records we tried to avoid the overlap in species names to make the study more accurate as possible, but since this work encompasses hundreds of species in different groups, including some species complexes (see supplementary material) some synonymy may occur.

## Results

Altogether, we found 18 orders of insects infected by Fungi and Oomycetes (58% of insect orders are infected). The great majority of fungal species are distributed among Ascomycota and Zygomycota, infecting 12 and 9 orders of insects respectively. Microsporidia species infect 12 orders, followed by Chytrids and Basidiomycota infecting 2 and Oomycota infecting just 1 order of insects. Below we examine in detail incidences of parasitism for each of these six groups focusing on the morphological and ecological traits that they evolved to make them efficient and specialized parasites of insects.

## 1. Incidence of entomopathogeneses

### 1.1 Oomycetes

The Oomycetes entomopathogens comprises 11 species spread among 6 genera, *Lagenidium* (1 species – *L. giganteum*), *Leptolegnia* (2 species – *L. caudata* and *L. chapmanii*), *Pythium* (3 species – *P. carolinianum*, *P. sierrensis* and *P. flevoense*); *Crypticola* (1 species – *C. clavulifera*); *Couchia* (*C. amphora*, *C. linnophila* and *C. circumplexa*) and *Aphanomyces* (1 species – *A. laevis*). They are known to infect just one order of insects, Diptera (Figure 1).

These insect pathogens can be found parasitizing their hosts in freshwater, primarily in well-aerated streams, rivers, ponds, lakes (Alexopoulos, 1996) or even treeholes (Washburn *et al.* 1988) and leaf axiles (Frances *et al.* 1989). Among them, the most well-known and broadly studied species is *Lagenidium giganteum* Couch, a facultative mosquito larvae parasite (Scholte *et al.* 2004). In an experimental study by Golkar *et al.* (1993), they describe the swimming behavior of zoospores towards the surface of the water. Their results, suggested that this behavior seems to be due to the cell shape and location of center of gravity rather than the result of a response given by a sensory system. However, chemotaxis of swimming spores has been described in some species (Cerenius & Soderhall, 1984).

The other genera like *Leptolegnia*, *Pythium*, *Crypticola* and *Aphanomyces*, have received only limited attention as entomopathogens (Scholte *et al.* 2004). Within genus *Leptolegnia*, only *L. caudata* de Bary (Bisht *et al.* 1996) and *L. chapmanii* Seymour (Seymour, 1984) have been isolated from insects (Scholte *et al.* 2004). The life cycle of *L. chapmanii* was presented by Zattau and McInnis (1987), reported infecting *Aedes aegypti*, the yellow fever and dengue disease vector. Effects of *L. chapmanii* on other aquatic invertebrates such as Odonata, Thichoptera, Coleoptera, Plecoptera and Cladocera were tested, though no infections were observed, suggesting high specificity (McInnis and Schimmel 1985). Members of the genus *Pythium* are species spread worldwide, occurring mostly as soil-inhabiting organisms or plant pathogens (Alexopoulos, 1996), whereas 3 species are known to infect insect larvae and even chronic skin lesions in mammals (Phillips *et al*. 2007). The main studies about this group are based on plant pathogen species, being poorly documented as entomopathogens. For more detailed informations about the entomopathogenic Oomycetes see Tiffney (1939), Su *et al.* (2001), Frances *et al.* (1989), Dick (1998), Seymour and Briggs (1985), Scholte *et al.* (2004).

### 1.2 Microsporidia

These are common pathogens of insects, being known 69 genera as type-host (compiled in Becnel & Andreadis, 1999) attacking 12 orders of these arthropods (figure 1). However, it is believed that all orders of insects posses Microsporidia among their pathogens. According to the same authors, the majority of infections are found in Diptera, with pathogenic species distributed among 42 genera. Ephemeroptera and Coleoptera, both with 5 genera infecting each, Lepidoptera with 4 genera, followed by Trichoptera with 3, Orthoptera, Odonata and Siphonaptera with 2 genera each and Thysanura, Hymenoptera and Isoptera with a single genus infecting them. These acounts are data based on genera which posse an insect as type-host, in this way this number tends to increase in future publications since an updated list will be published soon showing 93 genera of Microsporidians infecting insects (Dr. James Becnel, personal communication).

As will be discussed further, the Dipterans are the only group to present pathogens distributed among 5 of 6 groups of Fungi/Oomycetes (just lacking Basidiomycota infections, see figure 1), being the most susceptible order to fungal-oomycetes infections. Furthermore, this pattern remains among Microsporidian-diptera relationships. Amongst the 42 Microsporidians genera attacking Diptera, the largest, widespread and common group to infect these insects is the genus *Amblyospora* (Andreadis, 1994), attacking 79 species in 8 genera (Becnel & Andreadis, 1999). This genus posse a complex life cycle, which requires an intermediate copepod host and two mosquito generations in order to complete its full life span (Sweeney & Becnel, 1991).

Another important group among the entomopathogenic Microsporidia is the genus *Nosema*. They are considered for some authors the most important and widely distributed Microsporidia group (Tsai *et al.* 2003) being responsible for the majority of infections in Lepidoptera species (Zhu *et al.* 2003). A good example of their ecological and economical importance occur with species infecting bees (especially *N. bombycis* and *N. ceranae*), which are known to be responsible for great losses in apiculture, shortening bees’ life span (Higes *et al.* 2006). These infections are restricted to the midgut epithelial cells of bees and the infection occur by ingestion of spores. Once in the midgut, the spores are chemically stimulated to trigger the polar tube, which penetrates the host’s cells, starting the infection processes (de Graaf *et al.* 1994a). Later on, the spores are released with the feces, which are well adapted for survival in the environment due the presence of a characteristic thick walled tree-layered spore (de Graaf *et al.* 1994b; Vávra & Larsson, 1999).

### 1.3 Chytridiomycota

Among the entomopathogenic chytrids, we found 3 genera: *Myiophagus* (one species – *Myiophagus* cf. *ucrainicus*), *Coelomycidium* (one species – *Coelomycidium simulii*) and the most diverse genus *Coelomomyces* (63 species). Near all chytridiosis in insects were described in Diptera, however few examples of other infected groups were also reported. For instance, *Myiophagus* infecting scale insects (Muma & Clancy, 1931; Karling, 1948) in addition to dipteran pupae (Petch 1948). *Catenaria* is a genus of chytrids known as a parasite of cyst-nematodes, however, Doberski & Tribe (1978), found *Catenaria auxiliaris*, infecting Coleopteran larvae. Although, they are not sure if the colonization occurred after the larvae’s death (saprophytism) or if, in fact a parasitism relation occurred, leading the larvae to death. Thus, since this relationship is not proven yet, we will not consider *Catenaria auxiliaris* among the chytrids parasite of insects. The genus *Coelomycidium* is known to attack a specific group within Diptera order, the black flies (Jitklang *et al.* 2012; McCreadie & Adler, 1999). This disease is promptly identified by the observation of the larvae filled with spherical sporangia throughout the body cavity (Kim, 2011).

The *Coelomomyces* species are relatively well known and there is a significant literature about them, probably because their hosts are important human disease vectors (the mosquitoes: *Anopheles*, *Culex*, *Aedes* and *Simulium*) (Couch & Bland, 1985, pg. 285). Species within this genus can infect eggs (Martin, 1978), larvae (the great majority of infections)(see details in Travland, 1978) and also adults (Lucarotti, 1987). In some occasions, the fungus does not kill the larvae with its infection. When it occurs, the chytrid pass through the larval and pupal stage inside the host, reaching the adult stage, a mosquito. The preferential region of infection in adults is the ovary (Lucarotti, 1992). Once there, after a first blood meal the hypha matures to become sporangia, which is the fungal structure responsible to produce the zoospores (Lucarotti & Shoulkamy, 2000). Thus, instead of laying eggs, the mosquito will lay sporangia full of zoospores, ready to infect new larvae (Lucarotti & Klein, 1987).

### 1.4 Zygomycota

The Zygomycetes, as this group is commonly called, comprises a huge number of entomopathogenic species. These species are distributed among 20 genera: *Entomophthora, Conidiobolus, Entomophaga, Erynia, Massospora, Meristacrum, Neozygites, Sporodiniella, Strongwellsea, Pandora, Eryniopsis, Batkoa, Tarichium, Completoria, Ballocephala, Zygnemomyces, Ancylistes, Macrobiotophthora, Thaxterosporium and Basidiobolus*. It is difficult to say how many species of entomopathogens they are since many of these genera also infect different groups of hosts and due the constant taxonomic changes. However, since the scope of this work is not to provide a complete list of species of each group, but to present the diversity of morphology and strategies to infect their hosts, we will provide a broad figure of entomopathogenic species among Zygomycetes presenting some relevant examples of their diversity.

The Mucorales, the largest and morphologically diverse order within the class just present a single entomopathogen genus, *Sporodiniella* found attacking the Homopteran genus *Umbonia* in Ecuador (Evans & Samson 1977; Samson *et al.* 1988) and the Lepidopteran genus *Acraea*, in Taiwan (Chien & Hwang, 1997). But, by far the most important group of entomopathogens within zygomycetes is the Enthomophthorales.

The Entomophthoraceous fungi are well known as insect pathogens. Although the group comprises mostly insect pathogens species, some saprobes and specialized phytopathogens are also known, such as parasites of desmid algae and fern gametophyte (Humber 2012). While others parasitize mites, nematodes, and other invertebrate animals. This group attacks mainly adult insects, although two species of *Entomophthora* (*E. aquatica* and *E. conglomerata*) and *Erynia aquatica* are known to infect aquatic larval stages of mosquitoes (Scholte *et al.* 2004).

Regarding the transmission, according to Humber (1981), there are three distinct mechanisms for the forcible discharge of spores into the environment for the group (with exception of one single genus, *Massospora*, that release the spores passively, presented below). This study also suggests that this active discharge is a major event in the group’s life history, since it requires the expression of considerable portion of the genome in this crucial moment of the life cycle. Once the spore reach the aim host’s cuticle, the invasion of host body occurs by both mechanical and enzymatic ways (Brobyn & Wilding, 1977; 1983). Later on, after the fungal invasion, it kills the hosts by proliferation of mycelium or yeast-like cells and the production of secondary metabolites inside the insect body (Roy *et al.* 2006). After its death, the parasite stops the somatic growth (or decrease it highly), thus, growing exclusively reproductive structures (Alexopoulos, 1996).

These Fungi are characteristically biotrophics, consuming the host when they are still alive. This is one of the major differences if compared with Hypocrealean Fungi (discussed below in Ascomycota section), which are all hemibiotrophic, switching from a biotrophic phase, or parasitism, to a saprophytic phase, colonizing the host’s body even after its death (Roy *et al.* 2005). However, a bizarre phenomenon can also be observed in the entomophthoralean genera *Strongwellsea*, *Massospora*, and certain species of *Entomophthora, Erynia* and *Entomophaga*, (in addition, to the Ascomycete *Lecanicillium longisporum*) which produce the conidia (asexual spore) before the hosts death, in or on their living bodies (Roy *et al.* 2006).

The infections caused by *Strongwellsea castrans* in *Hylemya brassicae* and *H. platura* (Diptera) are classic examples of these peculiar situations (Nair & McEwen, 1973). In this case the infected fly is characterized by the presence of a large circular hole on the ventral side of the abdomen. But surprisingly, the infected insects can be observed acting normally, despite the big hole in its body, filled with fungal tissue and conidiophores (spore producing cells). Both, males and females were found infected, causing them castration and premature death (Nair & McEwen, 1973). Another similar case occurs with *Massospora cicadina*, which attack cicadas. This fungus also starts its sporulation still when the host is alive, walking and flying. The infected cicada will present the abdomen filled with fungal spores and the pressure due the swelling mass of fungus cause the continuing collapse of its body. Since the fungus maintain its growing inside the insect, time by time the host’s body keep collapsing and falling apart until remain just the head and thorax, exposing the fertile part of the fungus, thus disseminating the spores through the environment (Goldstein, 1929).

In response to these fungal infections, some insects can present abnormal activities such as unusual egg-laying behavior, leading to a decrease of egg hatching success, or even the female castration (Eilenberg, 1987). After fungal colonization in their bodies, some hosts were reported attempting to heal themselves by induced fever. Examples of this behavior are the flies infected by *Entomophthora muscae* and *E. schizophorae* (Roy et al. 2006; Kalsbeek *et al.* 2001), although this phenomenon was also described for another fungal groups and hosts (Blanford & Thomas, 1999). This behavior can play an important role in the insect’s population’s maintenance. According to Elliot *et al.* (2002), in fungal-infected locusts, those that achieved fever temperatures did molt, mature and reproduce, differently than that ones didn’t behave this way and eventually died quickly, demonstrating a clear benefit from the mentioned behavior.

## Basidiomycota

Although the Phylum comprises a huge amount of species (over to 22.000 teleomorph species, Hawksworth 1997), just 3 genera are known to infect insects. Those are *Fibulorhizoctonia* (species not formally described yet – see Yashiro & Matsuura 2007) infecting termite eggs, *Uredinella* (2 species – *U. coccidiophaga* and *U. spinulosa* infecting scale insects) and the biggest and most representative genus *Septobasidium* (ca. 240 species attacking scale insects – Hemiptera).

The Septobasidiales (i.e. *Uredinella* and *Septobasidium*, which the differences were discussed before) presents a peculiar and complex relationship with theirs hosts, the Diaspididae (Hemiptera). However, to understand its fascinating relationship we should first, understand even a little, about this insect’s biology: The Diaspididae comprise a group of small and sedentary insects, which spend their whole lives in one spot in a plant. Once a juvenile scale insect has found and established union with the host plant, it will stay there for the rest of its life, connected intimately with the plant by its long, slender sucking tube, feeding from the plant’s vessels. Since they are not able to defend themselves or fly away from enemies (see Heimpel & Rosenheim 1995), they do not survive unprotected (Christensen, 1975). These insects achieve this protection by secreting fine threads of white wax, which after 24h after their birth has formed a complete covering over the insect’s body. But, the challenges that this tiny insect should face in their lives are not restricted just by predation. Without proper protection they may perish with inclement weather forces such as dried up by the sun, washed off by rain, etc. Thus, the protection that the Septobasidiales fungus provides is very useful at colony level, since the fungus does not infect all of the individuals. Such sheltering is due to its abnormal growth pattern between layers, creating an elaborate system of tunnels and chambers inside its “body”, used by the insects as protection chambers for their whole lives (Couch, 1938).

On the other hand, there are also negative effects on insects such as dwarfism and incapacity of reproduction. The atrophy is probably due a result of haustoria activity, which is found attached only in the circulatory system or haemocoel. This specialized structure drains sap and nutrients from insect’s body, resulting in undernourishment. The content drained from the insect’s body was suggested by Couch (1931) to be stored in the conspicuous oil globules distributed along the fungus body. As the insects are kept inside the fungus, surrounded and held by hyphal threads, they become unable to move upwards and mate (Couch 1931, 1938). So, how they become infected?

The story begins few minutes after hatching, when the juvenile (crawlers) starts to seek out for a place to settle down (Couch, 1931). They can go through three different ways: 1) the first possibility is to stay on the same chamber with its mother, staying uninfected. 2) Others can move out from mother’s chamber to another one or even to another colony, at the risk to become infected by a spore on this way out. This group is also responsible to keep the colony growth, if the infected juvenile remains on its original colony (Couch, 1938). 3) The third group can settle down on an exposed spot, out of the colony, for example a bark surface. However, if juveniles on this third group become infected on their way, they will initiate a new fungus-insect colony. Nevertheless, the uninfected ones that follow this last option will be unprotected from extremes of weather, attacks of birds, predatory insects or from your main enemy, parasite wasps who lay the eggs inside the scale insect’s body (Evans, 1989). But, the fungus does not produce spores all over the year, giving some chance for many juveniles to be free from fungal infections (Couch, 1938).

The fungal spore producing is stimulated by warmth and moisture, throughout the spring and part of the summer, following rains. No basidiospore producing was described during the dry season. This reflects directly into the young infection rates, where, 1% is infected during dry season versus 50% infected during the wet season. Although there is no basidiospore formation, resting spores were observed being produced during the winter months, near the upper surface of the fungus body (Couch, 1931).

Regarding the less representative genus among Septobasidiales, the *Uredinella*, there are only two described species: *U. coccidiophaga* and *U. spinulosa* (see Couch, 1937; Couch & Petch 1941). They can be divided based on the substratum they are found infecting the insects, leaf and trunk to *U. spinulosa* and just trunk to *U. coccidiophaga*, and also based on teleutospores shape. As the genus infect a single insect, after the host’s death the fungus also die after producing spores in the Spring and reach sizes around 0.2 to 1.5 mm in diameter (Couch, 1937). On the other hand, *Septobasidium* have an undefined lifetime, since its body is “renewed” each season, when new infected insects remains at its original colony.

Another case of Basidiomycota parasitic of insects can be found between a *Fibulorhizoctonia* sp. (anamorph of *Athelia* – Atheliales) and some species of *Reticulitermes* subterranean termites. These termite workers are known to keep their eggs inside their nest in piles, thus taking care of them. However, Matsuura *et al.* (2000) found these piles, some sclerotia (globose fungal structures) being cared by the workers, as if they were eggs. The same study found that these sclerotia mimic the egg diameter and texture, thus the workers mistook the sclerotium for an egg.

Since the fungus life cycle is not clear yet, is difficult to say how the dispersion occurs. However, the same authors suggest two hypothesis: 1) the dispersion can occur by the alates carrying spores during swarming, although no spore formation were observed so far; 2) the fungi that form these sclerotia are ubiquitous and some workers eventually could bring them to inside the nest and once there it will pass as a true egg due its similarity to the eggs, even at microscopic levels (Matsuura et al. 2000). Even termites with no natural association with the “termite-ball”, when experimentally exposed to the fungus also presented the behavior to gather them together with their own eggs. Although there are just rare observations of this fungus actually infecting termite eggs, it occurs at natural conditions in Japan and USA (Matsuura, 2005) being considered to be facultative parasites of termite eggs.

For more details and species descriptions see: Couch 1931; 1938; Coles & Talbot 1977; Matsuura 2006; Yashiro & Matsuura 2007; Matsuura et al. 2009; Matsuura & Yashiro 2010; Lu & Guo 2010; 2011; Chen & Guo 2011a; 2011b).

## Ascomycota

As mentioned before, the phylum comprises since small groups of entomopathogens such as Pleosporales, Myriangiales, Ascosphaerales to big groups of the relatively better-known species like Hypocreales. Within the Pleosporales, the entomopathogenic species belong to the genus *Podonectria* (Petch, 1921), presenting unusual aspects among its class, like color and fleshiness. Due to these and other species’ traits, some researchers included them among the Hypocreales (Kobayasi & Shimizu, 1977) until their true nature was discovered (Alexopoulos, 1996). All known species were found infecting scale insects, covering the whole surface of insect body with cotton-like crust on which the perithecia will be produced and later, the multi-septated spores, not disarticulating into part-spores (Kobayasi & Shimizu, 1977). The genus is wordwide distributed, especially in warm climates (Petch, 1921) and the anamorphs related are the sporodochial genus *Tetracrium* (Petch, 1921; Kobayasi & Shimizu, 1977) and the pycnidial genus *Tetranacrium* (Rossman, 1978; Roberts & Humber 1981).

The Myriangiales includes a number of species associated with plants, resins, or scale insects on plants (Alexopoulos, 1996). The entomopathogen species present perennial growth for several years, or at least until the branch, where the scale insect is attached, dies. The hosts can be found directly under each stroma, where there are several scales, penetrated and covered by mycelium (Miller, 1940). The stroma sometimes can be formed at the side of the insect (Petch, 1924). The growth occurs when the rains begin, where the live parts of the fungus body increase in diameter, producing ascocarps and later on ascospores. Reproduction is entirely by ascospores and no evidence of conidial (asexual spore producing cells) formation was found on stromata of any age neither in culture (Miller, 1938). For taxonomic and additional discussions see Petch (1924) and Miller (1938; 1940).

A unique group of bee pathogens are also included in this Phylum. The genus *Ascosphaera* is a group of approx. 26 species, which are specialists in exploit bee’s provisions. They are mostly exclusively saprophytes on honey, cocoons, larval feces or nest materials such as leaf, mud or wax of bees (Wynns et al. 2012). However, some species are known as widespread fungal disease agents, attacking broods of numerous species of solitary and social bees, in a disease called chalkbrood (Klinger et al. 2013). The infection occurs when the larva ingest fungal spores, being killed by the hyphal growth inside its body. Then, leading to new spore formation on the cuticle of the dead larvae (Vojvodic et al. 2012). The morphology of *Ascosphaera* is very peculiar when compared to other fungal groups. The ascoma is a small brown to blackish brown spore cyst, a single enlarged cell containing ascospores (Wynns et al. 2012). In addition, they also lack a well-developed ascocarp, which led them to be considered as yeasts (Alexopoulos, 1996). For a detailed life cycle see McManus & Youssef (1984).

The Hypocrealean fungus encompasses important genera of entomopathogens Fungi such as *Hypocrella, Cordyceps, Ophiocordyceps, Moelleriella* among others. In addition to these species, there are a myriad of anamorphic species related to them, i.e. *Hirsutella, Metarhizium, Hymenostilbe, Akanthomyces*, etc (Roberts & Humber, 1981). In the past, these anamorphic species (which only the asexual stage is known) were treated traditionally as a separate group, the Deuteromycetes. But, nowadays with the advancing molecular studies, some of these species are now strongly supported to be asexual stages of mainly Ascomycota (see Liu et al. 2000). Even 25 years ago, Evans (1988 book) included both in the same section in his book’s chapter and said: “*The entomopathogenic Deuteromycotina are included within this subdivision* (Ascomycotina section of his book) *since many of them are either proven or purpoted anamorphs of Ascomycotina*”. Here, since many anamorphic species were proven to be part of life cycle of ascomycete teleomorph phase, we will just address the teleomorph names to avoid some synonymy and to follow the new “One fungus one name” code.

Within this largest group, the Hypocreales, we can highlight some genera. For example *Hypocrella*, which was beautifully monographed by Chaverri *et al.* (2008). These Fungi are known to infect white flies and scale insect in tropical forests, with few species recorded to sub-tropics. They possess high virulence causing epizootic in their host’s population. Not just these cited species are able to cause epizootics, also species of *Ophiocordyceps*, specially attacking ants, are known to cause huge infestations in small areas, those called graveyards (Pontoppidan *et al.* 2009; Evans & Samson 1982; 1984). Indeed, one of the most fascinating phenomenons regarding entomopathogenic Fungi is the one caused by *Ophiocordyceps* on ants, originally described by Tulasne & Tulasne (1865) as a *Torrubia* species. They infect the ant’s brain, leading them to the death, by changing its behavior in a way to lead the insect to reach an optimum microclimate site required by the Fungi (see the complete story in Andersen et al. 2009). After infection, the parasitized ant, so-called zombie-ants (Andersen & Hughes 2012), leave the colony to die biting tightly on abaxial or edge of the leaf, twigs, branches, etc (the death position is related to each species). Later on, after ant’s death, the fungus start to grow hyphae to glue them to the leaf, avoiding the ant’s cadaver to fall on the ground, precluding the fungal growth, spore dispersion and further transmission to new hosts.

3.0 Diversity of exploitation strategies cuckoo, living insects

## Discussion

### Factors promoting diversity

Among the entomopathogens addressed in this paper we can highlight some ecological groups. For instance, the hyper diverse pathogens of sap-sucking Hemiptera, one of the most diverse groups of entomopathogenic organisms. Based on counts of teleomorphic names of Hypocreales (Ascomycota) we found 181 species of fungi parasitizing exclusively Hemipteran insects (figure 2). Most cases were infections of adult stages (n=166 records). The *Hypocrella-Aschersonia* species are responsible for the majority of infections, with approximately 95 species infecting scale insects (Coccidae and Lecaniidae, Hemiptera) and the white-flies (Aleyrodidae, Hemiptera). The other infections caused by Hypocreales in Hemipterans are spread mainly among the genera: *Moelleriella* (23), *Ophiocordyceps* (19), *Torrubiella* (18), *Cordyceps* (17) and *Samuelsia* (6). In addition, another groups such as Myriangiales (Ascomycota), Septobasidiales (approx. 240 species - Basidiomycota) and even the chytrid fungus *Myiophagus* (Chytridiomycota) are known to infect the hemipteran sap sucking insects.

**Figure 2:**
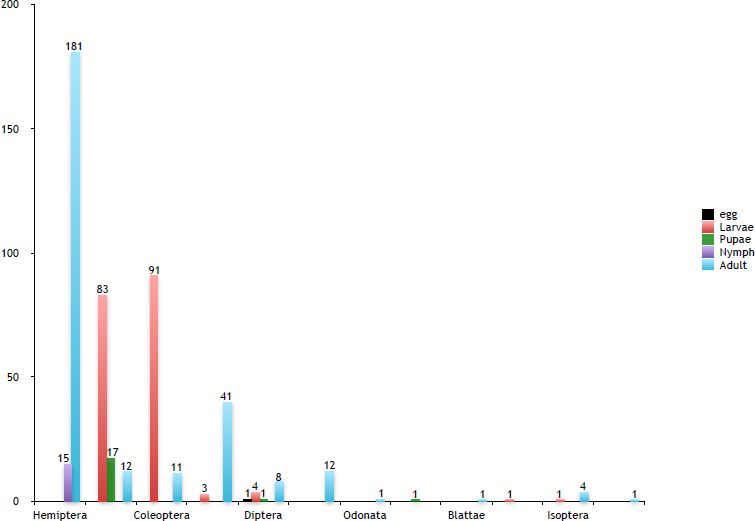
Hypocreales teleomorph species numbers and distribution on each stage of host’s development.

**Figure 3:**
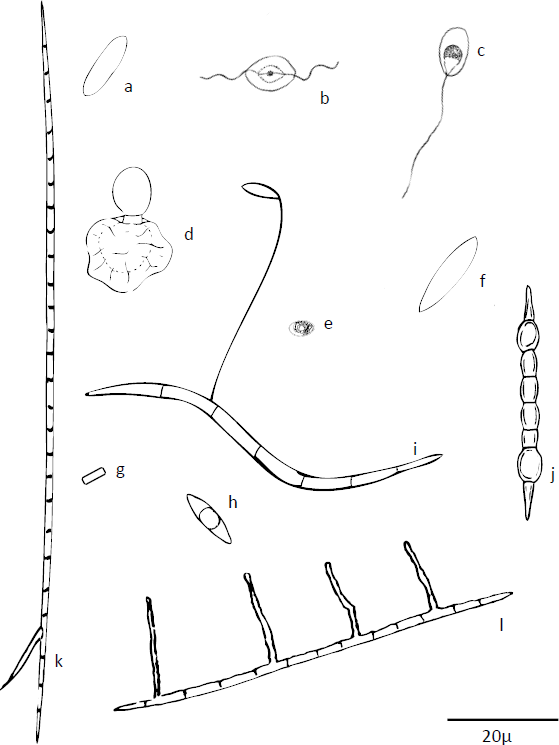
Spore diversity within entomopathogens Fungi (in μ): a – *Moelleriella sloaneae* (13-15 × 2.8-3) – Chaverri *et al.* 2008 b – *Lagenidium giganteum* (8-9 × 9-10) – Couch, 1935 c – *Coelomomyce spsophome* (5 × 10) – Whisler *et al.* 1975 d – *Entomophthora thripidum* (10-15 × 8-12) – Samson *et al.* 1978 e – *Nosema hyperae* (3.1 × 1.7) Ovcharenko *et al.* 2013 f – *Septobasidium maesae* (18-19.5 × 4-5) Lu & Guo, 2009 g – *Ophiocordyceps lloydii* (4 × 1) - Kobayasi & Shimizu, 1978 h – *Hypocrella raciborskii* (10-16 × 2.5-4) – Luangsa-ard*et al.* 2007 i – *Ophiocordyceps camponoti-rufipedis* (80-95 × 2-3) Evans *et al.* 2011 j – *Ophiocordyceps blattae* (40-60 × 4-6) – Luangsa-ard *et al.* 2010 k – Ophiocordyceps camponoti-melanotici (170-210 × 4-5) – Evans *et al.* 2011 1 – Ophiocordyceps camponoti-novogranadensis (75-95 × 2.5-3.5) – Evans *et al.* 2011

However, to understand such wide range of entomopathogen species infecting hemipterans, we need to even superficially understand the origin and evolution of its dominant pathogens, placed in Hypocreales order. According to Spatafora *et al.* (2007) and Sung *et al.* (2007; 2008), the entomopathogenic Hypocreales had their origin from Fungi associated with plants, although some modern groups remains associated with them (i.e. Bionectriaceae). Throughout the course of evolution, were estimated around 5 to 8 host-shifting events (Spatafora *et al.* 2007). For instance, from plant-pathogen fungus to animal parasites, from animal pathogens to plant endophytes, from animal pathogens to hypogeous fungus pathogens, etc (Spatafora *et al.* 2007; Nikoh & Fukatsu 2000). Therefore, these successions of shifting along the evolutionary history of the group shaped the diversity we have currently, with species attacking fungi, plants and invertebrates, mostly insects, in the same clade.

But, why Hemipteran-Fungi is so taxonomically diverse and how the Fungi reached this group of insects? Reconstructing the modern insects evolution, we can clearly see a great diversification observed by the completely different morphologies shaped by evolutionary processes. We can highlight the mouthparts as one of the most important characters to recognize how certain groups of insects feed, exploit and interact with the environment. In the case of Hemipterans, their mouthparts evolved into two pairs of long and fine stylets, being able to create strong suction in order to drawn fluids from plant tissue (Grimaldi & Engel, 2005). This derived feature was essential for this group of insects, since it allowed them to exploit a new niche: living on the plants and feed from their sap. However, in the same way that insects evolutionarily adapted its mouthparts into stylets in order to exploit another environment, the Fungi in order to exploit another food source would also switched from plant to insect body. An indication that these Fungi achieved such a great success on this host shifting can be observed in the species-richness we have currently on tropical forests worldwide.

But, one question arises: How did the fungus switched from plant to insects? Observing the figure 4, we can see a clear relationship between taxonomically distant groups: truffles (Fungi), grasses (Plantae) and Hemipterans (Animal). This co-occurrence in the same habitat considerably increases the chance of encounter of an endophytic grass fungus with the grass’ exploiter sap-sucking Hemiptera, due obvious reasons. So, applying this scenario to the “Host habitat hypothesis” which says that: “*host-shifts should follow micro-habitat or feeding habit lines, on the grounds that probability of encounter is a dominant factor in the evolution of host associations* (Shaw, 1988), we can suggest that this fungal shift from endophytes to entomopathogens was due this close relationships between both hosts: plant and insect. In support to this idea, we can observe a close phylogenetic relationship between species of fungi with completely different hosts (different kingdoms) that share very close niches or “feeding-habit” (Figure 4). Afterwards, reaching and (eventually) successfully colonizing the new insect host, the fungi was gradually adapting to its ecology, in order to optimize the horizontal transmission, clearly observed by the spore adaptations. For instance, the *Hypocrella-Aschersonia* clade, which presents mitotic slime spores, adapted for short distance dispersal by rain-splash on leaf surfaces, which is a hotspot to find these hemipterans (Evans, 1989; pg. 221). It is important to mention that *Hypocrella-Aschersonia* clade is the only entomopathogenic group within Hypocreales to present plant pathogens and endophytes as the sister group (figure 4), demonstrating the close phylogenetic relation between these two ecologically distinct lineages of fungi (see complete phylogeny of the group and its distribution of host-kingdoms in Spatafora *et al*, 2007).

**Figure 4:**
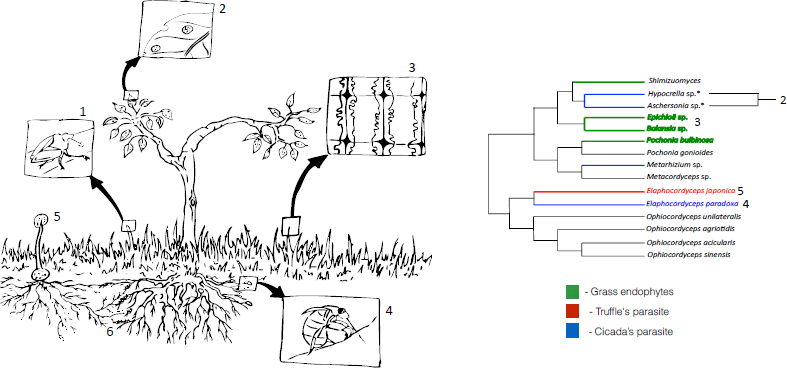
The close relationship between Hemipterans and entomopathogen Fungi. 1 – Hemiptera feeding from a plant; 2 – *Hypocrella-Aschersonia* entomopathogens; 3 – Fungus associated within the plant cells (endophyte); 4 – Hemipteran nymph feeding from plant roots (host of some species of *Elaphocordyceps* and *Cordyceps*); 5 – *Elaphocordyceps* infecting *Elaphomyces*; 6 – *Elaphomyces* in mycorrhyzal association. (Adapted from Spatafora *et al.* 2007)

**Figure 5:**
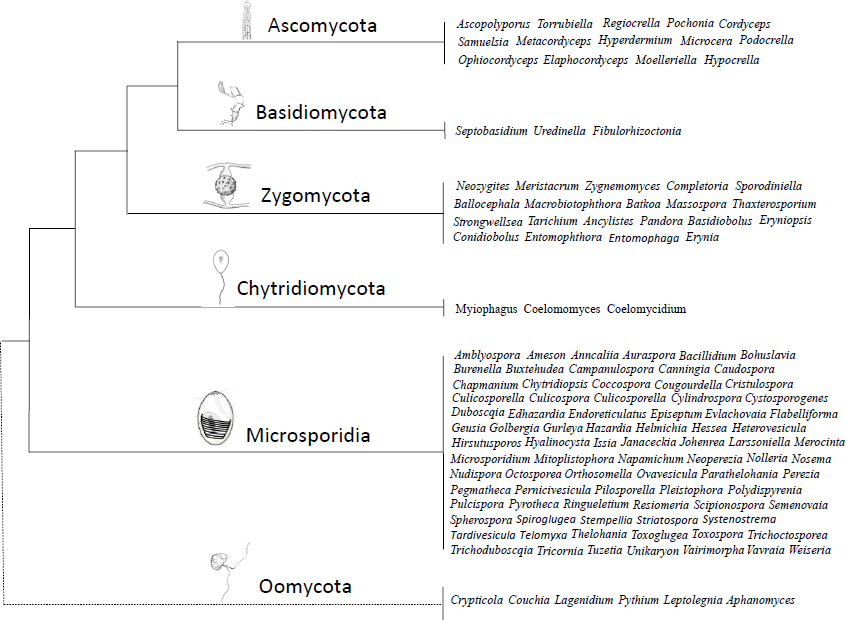
Diversity of genera of entomopathogens across fungal and Oomycets

## Broad range of ecologies, broad range of pathogens

The Dipteran was the only order with records for all stages of development, eggs, larvae, pupae and adults, infected by entomopathogen Fungi and Oomycetes and only Basidiomycota did not occur infecting them. It is probably because this is the most ecologically diverse group of insects, which is found all over the world, occupying a broad range of niches: blood feeders, endo- and ectoparasites of vertebrates, gall makers, larval and adult predators, leaf miners, parasitoids, pollinators, saprophages, wood borers, etc (Grimaldi & Engel, 2005). In addition, the fly’s larvae can be found in many different breeding sites such as aquatic, semi-aquatic (wet soils, stones at stream edges) or terrestrial (Fungi, rotten wood, trunk, etc). Moreover, the high diversity of ecologies among the different groups of pathogens infecting Dipterans can also support this hypothesis: the aquatic Oomycetes and chytrids and all fungal phyla with exception of the Basidiomycota, which are highly adapted to infect Hemipterans. Thus, we suggested that occupying such diverse niches increase the chance to get infected by fungal and oomycetes pathogens, which also occupy a myriad of environments.

Beyond the infected groups discussed above (Hemiptera and Diptera), we can also highlight the great prevalence of infections of larval stage of Lepidoptera and Coleoptera (Figure 2) by Hypocrealean fungi (Ascomycota). The larvae of both orders are the most common host for the two important genera of entomopathogenic Hypocreales, the *Cordyceps* and *Ophiocordyceps* (Figure 2). As in the case of Hemipteran infections, we cannot explain yet precisely the reason of the prevalence of entomopathogens infections on larval stage of Coleoptera and Lepidoptera orders.

However, we suggest the consideration and comparison of some biological traits between the larvae and adults, which can be crucial to understand such scenario: 1) Partition of niches, 2) Predictability in time-space scales, 3) Feeding rate and 4) Protection (Cuticle). (1) As being holometabolous insects (which posses complete metamorphosis), the larval and adult stages are ecologically separated, occupying completely different microenvironments, thus avoiding competition between juvenile and adults (Gullan & Cranston, 2010; pg. 234). (2) Both, Coleopteran and Lepidopteran larvae generally presents slightly mobility compared to the wandering adults (especially Lepidoptera), tending to be closer to the breeding site eating ferociously, thus being more predictable in time-space scales. (3) The larva needs to eat an impressive amount of food and store them, in order to grow as quick as possible, making them a huge reservoir of energy. (4) Furthermore, since larvae need to grow in high rates, it would be impossible if they had the hard exoskeleton (especially Coleoptera) adults have. However, on the other hand, having such soft and thin skin would make these organisms much more easier to be invaded by fungal spores equipped with their enzymatic and physical apparatus. It is worthy to say that the usual defenses the larvae usually present (mimicry, aposematism, gregarious behavior, stinging hairs, etc) that are very useful against predators are completely useless against fungal pathogens.

